# The roles of wrinkle structures in the veins of Asian Ladybird and bioinspiration

**DOI:** 10.1101/2020.01.02.893388

**Authors:** Zelai Song, Yongwei Yan, Wei Wu, Jin Tong, Jiyu Sun

## Abstract

The deployable hind wings of the Asian ladybird beetle (*Harmonia axyridis*) play important roles in their flight. Wrinkle structures of veins are found on the bending zones of the hind wings of *H. axyridis*. This paper investigates the effect of the wrinkle structures of the veins of the hind wing on its deformation. Based on the nanomechanical properties of the veins, morphology of the hind wing, surface structures of veins and microstructures of the cross sections, including the veins and wing membranes, we establish four three-dimensional coupling models for hind wings with/without wrinkles with different and uniform reduced modulus. Relative to the bending and twisting model shapes, Model I, which includes the wrinkle structure and different reduced-modulus veins, has much more flexibility of passive deformation to control wing deformations. The results show that both the wrinkle structures in the bending zone and varying reduced modulus of the veins contribute to the flight performance of bending and twisting deformations of the hind wings, which have important implications for the bionic design of the biomimetic deployable wing of micro air vehicles (MAVs).

## 1. Introduction

Insects provide the bioinspiration to design light-weight and miniature micro air vehicles (MAVs) (Takizawa et al., 2015; Phan et al., 2016; Rajabi et al., 2015). The Asian ladybird beetle, *Harmonia axyridis* can fly at a high velocity and has a strong propensity for long-distance dispersal (Fig. 1) (Maes et al., 2014; Jeffries et al., 2013); in particular, the deployable hind wings of *H. axyridis* can reduce its body size (the folding ratio exceeds 2) (Sun et al., 2018a). To reduce the size and increase the portability of wings for MAVs, the designs of bioinspired species with folding wings have more advantages (Truong et al., 2014; Jitsukawa et al., 2017; Faber et al., 2018).

**Fig. 1.**
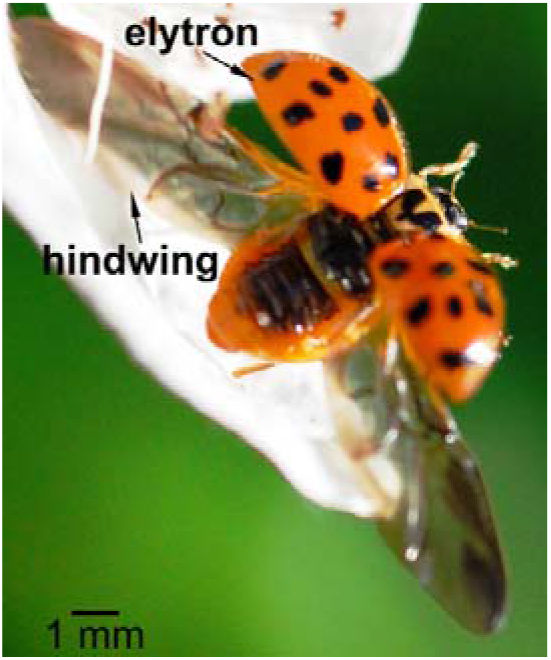
Ladybeetle taking off.

The biomechanical and flight performances are related to the microstructures of insect wings (Wang et al., 2017; Gorb, 1999). Insects can control their wings via active and passive deformations that result in a flexible flight performance (Rajabi et al., 2016; Jongerius and Lentink, 2010; Kesel et al., 1998; Mountcastle and Daniel, 2009). The timely deformations of wings can lead to high lift and improve the distribution of aerodynamic forces on the wing during flight (Hou et al., 2017). The flapping wings of insects are composed of wing membranes and veins (Bao et al., 2011; Schieber et al., 2017). The structure and patterns of the veins of insect wings affect their mechanical properties (Combes, 2003; Meyers et al., 2008). The structural constitution of the wing vein wall is characterized by gradual transition zones between the wing membrane and vein wall and an internal layer of elastic material (Bergmann et al., 2018). Some veins of the wing of insects are hollow tubes that allow hemolymph to flow (Zhao et al., 2013) and help to unfold the hind wings via a hydraulic mechanism (Sun et al., 2014). The anterior wing margin is sharply bent when folded (Haas and Beutel, 2001), so the bending zone of the hind wing is a significant part of the folding/unfolding mechanism (Haas, 2006).

The modulus of a vein varies in the wing at different positions (Ha et al., 2011; Herbert et al., 2000a), which is also related to the density (Bao et al., 2011), specific strength, toughness and hardness (Oliver and Pharr, 1992; Oliver and Pharr, 2004). The modulus and hardness are not unique among different beetles species, and the positions of sampling and the functional heterogeneity of biological samples are considered to result from biological evolution (Yu et al., 2013). The effects of blood in different veins with identical modulus on the vibration characteristics have been analyzed in the dragonfly wing (Hou et al., 2015). To understand the relationship of the structural properties and function, the wing and veins are used the same modulus for the simulation (Rajabi et al., 2017). In analyzing the effects of the camber and stress stiffening of the membrane, the model assumes that the veins have an identical modulus (Ha et al., 2013). Wing models of insects have also been used as foundations to simulate the mechanical properties of wings (Tong et al., 2015).

In this paper, 3-D structural models of the hind wing are first established based on its microstructures. Then, according the nanomechanical properties of the veins at different positions, four coupled models are established (models I and ◻: wrinkled veins with different reduced modulus and unique modulus, respectively; models ◻ and ◻: smooth veins with different reduced modulus and unique modulus, respectively). In the following sections, the effects of the structures and nanomechanical properties of the veins on bending and twisting deformations are discussed.

## 2. Materials and methods

### 2.1. Specimen

Specimens of *H. axyridis* (Polyphaga, Coccinellidae) are collected in Changchun, China in April 2019. Alcohol is used to anesthetize the specimens for the test. Thirty male *H. axyridis* with relatively similar sizes and elytra are selected for this study. Each specimen is measured 5 times. Small pieces of the hind wings are carefully removed from the bodies using sharp razor blades.

### 2.2. Microstructure

To obtain the entire hindwing morphology of *H. axyridis*, a stereomicroscope (OLYMPUS SZX7, Olympus Optical Co., Ltd., Tokyo, Japan) is used.

To investigate the microstructures of the veins, scanning electron microscopy (SEM) (Zeiss EVO18, Carl Zeiss AG, Germany) is used.

To obtain the diameter and thickness of the veins and membrane wing, hematoxylin-eosin staining and inverted fluorescence microscopy (OLYMPUS DP80, Olympus Optical Co., Ltd., Tokyo, Japan) are employed. The hind wings are kept in a unique stationary liquid for 24 hours that is composed of glacial acetic acid, ethyl alcohol and formalin. Then, the hind wings are dehydrated and cut into six pieces at the nanoindentation test points at the C+ScA after being embedded in paraffin using a paraffin embedding machine (Leica EG1150C, Leica Biosystems, Germany) to support the specimens in the slicing test. The paraffin slicing machine (Leica RM2235, Leica Biosystems, Germany) is used to cut each specimen into paraffin sections. To easily observe the cross section, HE staining is applied to the paraffin sections, which are covered by cover glasses.

### 2.3. Nanoindentation

To investigate the nanomechanical properties of the veins and membranes of the wings of *H. axyridis*, a nanoindenter (Nanaoindenter, Hystiron, Inc., USA) is used to test ten locations of the veins, as shown in Fig. 2A. The test locations have to be flat without microtrichia; otherwise, the test results will be affected. A Berkovich indenter tip is used (diamond, Young’s modulus: 1140 GPa; Poisson’s ratio: 0.07, with a half include angle of 65.27°). The load profile is consisted of three 10-s intervals: a linear increase from zero to the maximum load, a constant maximum load, and a linear decrease back to zero. A trapezoidal loading function is applied for the indentation tests. A peak load of 100 μN is applied to penetrate the articular surfaces of the membrane wing and veins over a range of depths (100–150 nm). Force-displacement curves are used to determine the effective reduced modulus (*E*_*eff*_) of the material surfaces (Oliver and Pharr, 1992). The results are found to be independent of the loading rate at approximately 50 μN/s. This analysis is based on the simple assumption that the unloading is fully reversible. The analysis is performed under ambient conditions.

**Fig. 2.**
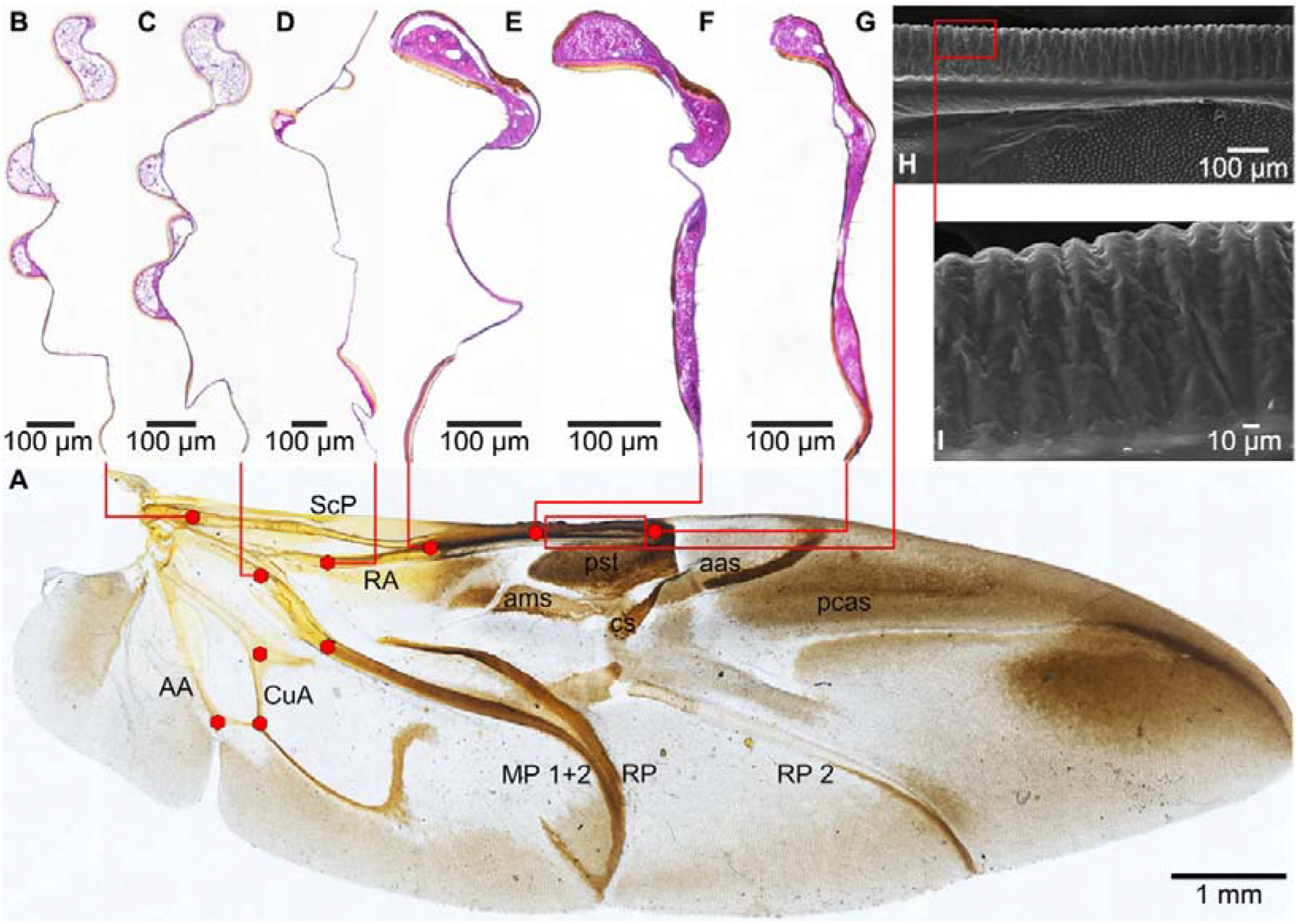
A *H. axyridis* hind wing and the nanoindentation test points. (B-G) Cross sections of the first six positions cut at the test points of the hind wing. (H and I) Surface structure of the vein from position F to G. ScP: Subcosta posterior. RA: Radius anterior. cb: Costal bar. ams: Anteromedian sclerotization. CuA: Cubitus anterior. AA: Anal veins. MP1+2: Media posterior1+2. RP: Radius posterior. arc.c.: Complex arculus. cs: Central sclerotization. pst: Pterostigma. aas: Antero-apical sclerotization. RP2: Radius posterior 2. pcas: Postcosto-apical sclerotization.

The Oliver-Pharr method has been accepted to be able to determine the Young’s modulus (*E*) and hardness (*H*) (Oliver and Pharr, 2004). The equations are as follows:

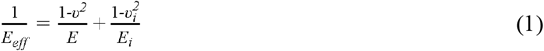

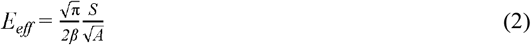

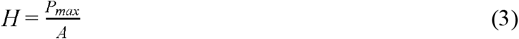

where *v* and *v*_*i*_ are the Poisson’s ratios of the specimen and indenter, respectively; *E* and *E*_*i*_ are the Young’s modulus of the specimen and indenter, respectively; *A* is the resultant projected contact area between the tip and the sample surface; *P*_*max*_ is the maximum indentation force; and β is 1.034 for the Berkovich tip (King, 1987), which is a constant related to the tip geometry. *S* = *dP/dh* is defined as the slope of the upper portion of the unloading curve during the initial stages of unloading and is independent of the work-hardening behavior of the veins. When *h*_*f*_/*h*_*max*_ < 0.7 (Bolshakov and Pharr, 1998; Pharr, 1998), where *h*_*max*_ is the indenter displacement at the peak load and *h*_*f*_ is the final depth of the contact impression after unloading. The function parameter of the simulation test is used as the reduced modulus.

### 2.4. Finite element modeling

ANSYS^®^ is used to calculate the bending and twisting deformation due to the same force. Then, we compared the reaction deformations of four hind wing models to select the best hind wing model.

#### 2.4.1. 3-D coupled models

Both the structures and mechanical properties of the hind wing determine how it deforms in response to applied forces. Thus, considering these two factors, 3-D coupled models are established to investigate their effect on wing deformation. The bending zone of the veins also has wrinkles (Haas et al., 2000), so the geometries of the hind wings are designed by the Solid Edge software, whose outline size and vein trace are followed by the real hind wing shown in Fig. 2. The diameter of each vein is assumed to be uniform from the base to the tip, and the veins are assumed to be smooth circular pipes. The material properties of veins are designed as the real circumstance. The membrane wings of the four models are assumed to have identical thicknesses and material properties and to be flat. The diameters of the veins (*R*) are assumed to be 0.047 mm, the inner diameters (*r*) are 0.023 mm, and the thickness of the wing membrane is assumed to be 0.01 mm. The wing models shown in Fig. 3 approach realism since they consist of a flat uniform membrane, a limited number of veins, and a uniform thickness.

**Fig. 3.**
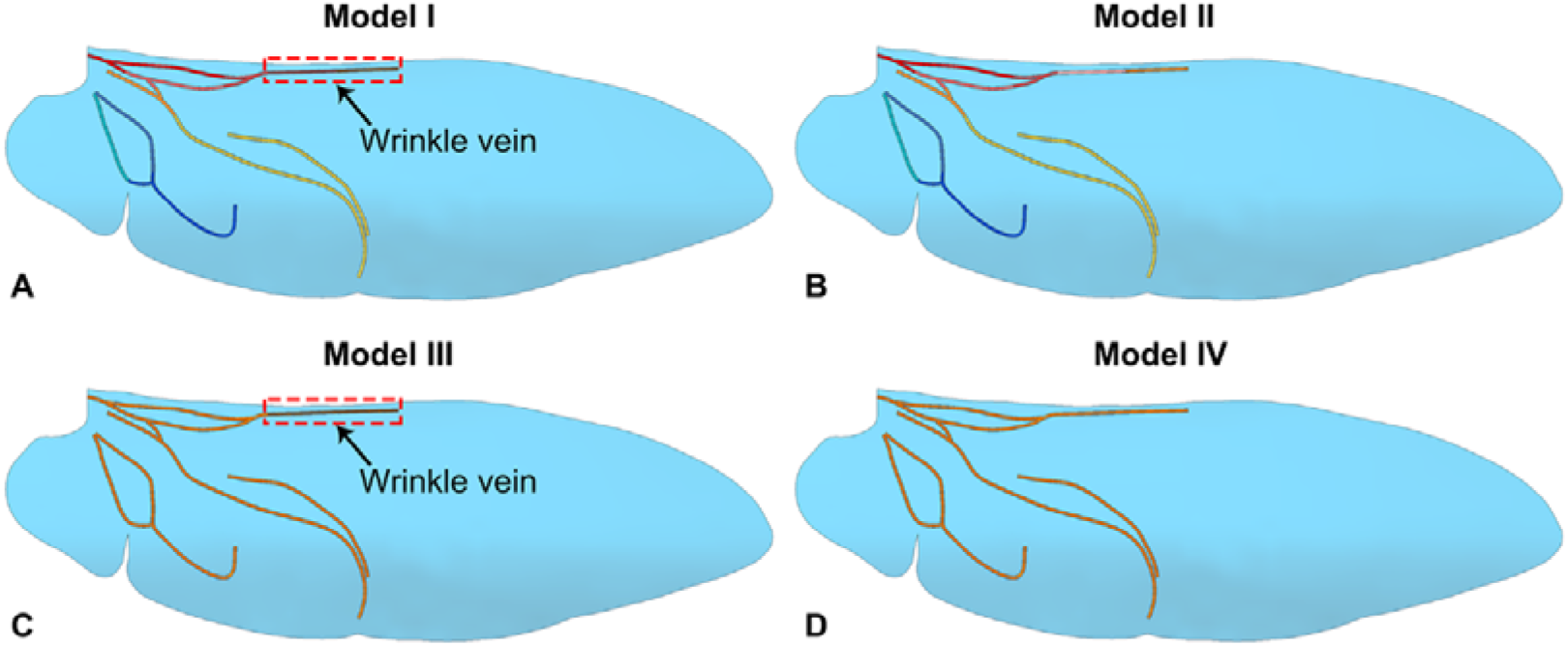
Models of the hind wings. (A) Model I with different reduced-modulus veins with wrinkles. (B) Model II with different reduced-modulus veins without wrinkles. (C) Model III with the uniform reduced-modulus veins with wrinkles. (D) Model IV with the uniform reduced-modulus veins and without wrinkles. The red dashed box zone is a vein with wrinkles.

As Fig. 4 shows, the vein models with/without wrinkles are designed using the SEM images shown in Fig. 2I, which are of the last half of the ScP. The structures of the veins of the four models are designed from two veins without/with wrinkles. The veins or structures indicated by the red dashed line are used to explore the function of veins with wrinkles.

**Fig. 4.**
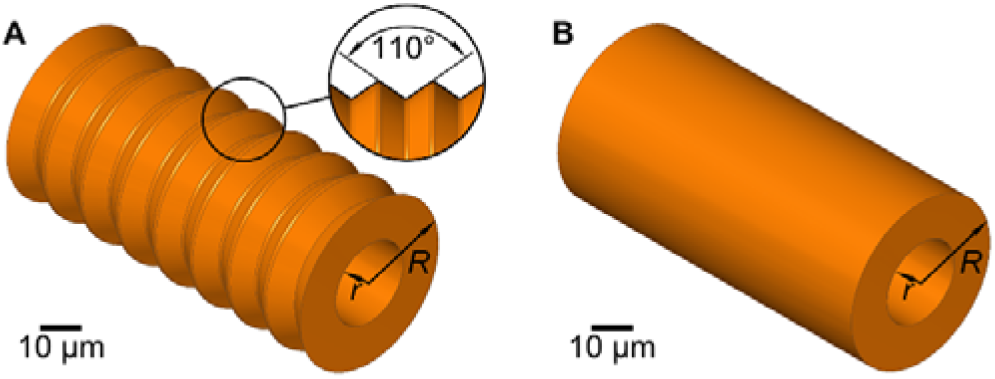
(A) Vein model with wrinkles. (B) Vein model without wrinkles. *R*: diameter of the veins; *r*: inner diameter of the veins.

#### 2.4.2. Mesh generation

A mesh convergence analysis is performed on the models with different numbers of elements to achieve a compromise between the accuracy of the results and the computational time. The wing membrane and veins are meshed using an automatic method due to the complex geometry. The mesh is refined to converge the solution. To determine the minimum number of elements necessary to capture the bending behavior of the wing, we meshed the models of the wings and veins with element sizes of 0.008 mm and 0.006 mm, respectively, and we found that the element size of 0.006 mm is sufficient to ensure the performance of the model. In total, approximately 1855750, 1856495, 2124544 and 2114650 elements are required for the four hind wing models with different reduced-modulus veins and the same-reduced modulus veins to obtain results that are independent on the mesh size. The hind wing models are mesh-independent for meshes with 7176268, 7176797, 7465001 and 7417545 nodes. The vein models with/without wrinkles are meshed using the automatic method with approximately 3485 and 2400 elements with 17320 and 11676 nodes, respectively.

#### 2.4.3. Loads setting

In this study, we only focused on comparing three wing models. Hence, the modulus of the membrane and veins does not affect the direction of deformation. Thus, the membrane materials are assumed to be isotropic and homogenous and to represent the average reduced modulus of the membrane.

The costal margin of the hind wing can rotate relative to the base of the leading edge vein (Betts, 1986a). The twisting force is assumed on the ScP and AA with the pressure, and the bend force is assumed on all of the veins with the pressure.

The hind wing models are fixed at the wing base with zero displacement and rotation. The bending and torsional moments induced by the aerodynamic forces during flight are predominantly carried by the longitudinal veins (Haas and Wootton, 1996). There is a main longitudinal vein in the hind wings of *H. axyridis* in our models. Therefore, assuming the same aerodynamic lift generated by the hind wings, we can expect that the part of hind wings modelled in our study is subjected to an aerodynamic force that is almost equal to the insect body weight (23 mg) (Yasuda and Ohnuma, 1999; Sun et al., 2018b; Jongerius and Lentink, 2010) during hovering flight. To analyze the effect of the veins on the deformation of the hind wing, the load is always set on different veins. The area of the hind wing is approximately 18.41 mm^2^, as shown in Fig. 2A. The equivalent pressure on the hind wing is 6.25×10^−6^ MPa, and the applied force is set on the end of the vein with wrinkle as 2.3×10^−4^ μN. As Fig. 5 shown, the schematic diagram shows the force application on hind wing for bending and twisting deformation.

**Fig. 5.**
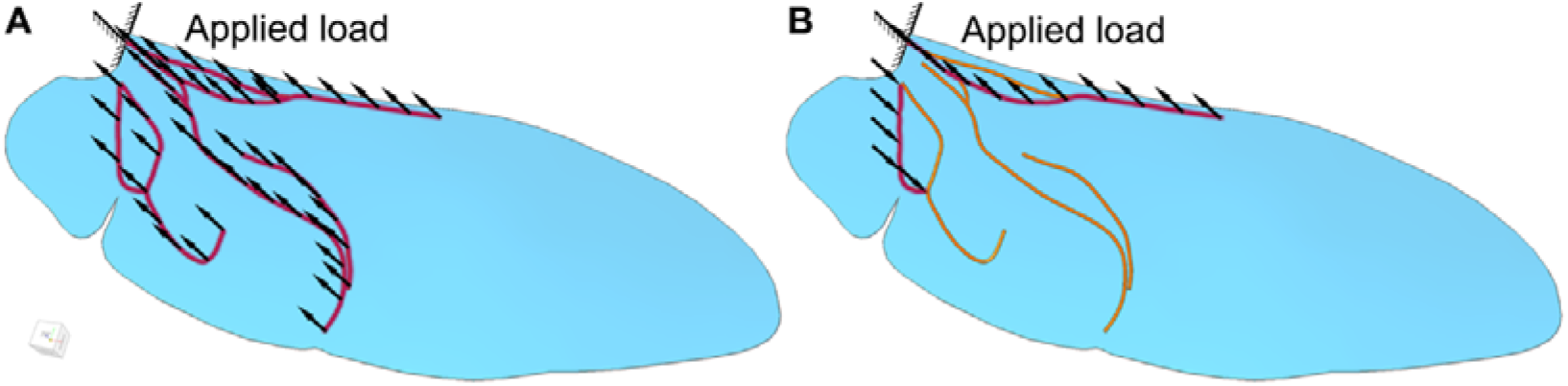
Schematic diagram of force application on bending deformation (A) and twisting deformation (B). The arrows indicate the direction of force application.

#### 2.4.4. Modal characterization of hind wing

To analyze the nanomechanical properties of the veins, the structure of the hind wing and veins are analyzed, and the reduced modulus is measured. Poisson’s ratio is assumed to be 0.25 (Saha and Nix, 2002; Bell et al., 1992), which is unknown. In formulas (1) and (2), the reduced modulus (*E*_*eff*_) of the specimen approached Young’s modulus (*E*).

## 3. Results

### 3.1. Microstructure of the hind wing

Fig. 2 shows the hind wing of *H. axyridis* and the cross section of the hind wing. The main veins are always on the first half of the hind wing, but the latter half of the hind wing, whose function is folding/unfolding, almost always includes a wing membrane and false veins (Schieber et al., 2017). The hind wing also consists of the epicuticle, mesocuticle, central lamella and exocuticle, and the epicuticle, which is dark purple, is also on the surface of the hind wing (Schieber et al., 2017). The cross sections of an artery and vein are composed of the endothelium, tunica intima, tunica media, and tunics adventitia(Meyers et al., 2008). The orange positions of the cross sections of the hind wing are the exocuticle, and the purple positions are the mesocuticle, as shown in Fig. 2.

### 3.3. Nanomechanical properties of the veins

To ensure the true nanomechanical properties of the veins, a flat plane is selected for the veins without microtrichia on the dorsal side of the hind wing because membrane wing is composed of the epicuticle, mesocuticle, central lamella and exocuticle, and the epicuticle (Schieber et al., 2017) and the veins are composed of the endothelium, tunica intima, tunica media, and tunics adventitia (Meyers et al., 2008), so the cross section of the hind wing cannot be used to measure the combination property of the hind wing.

The mean reduced modulus (*E*) and hardnesses (*H*) of ten zones of the hind wing are shown in Fig. 6. The reduced modulus widely varies with the vein location. In these ten zones, *E* is 2.29± 0.18 GPa, 3.12± 0.25 GPa, 5.81± 0.40 GPa, 8.40± 0.21 GPa, 10.06± 0.59 GPa, 2.61± 0.32 GPa, 2.00± 0.12 GPa, 7.78± 0.48 GPa, 3.08± 0.17 GPa and 6.52± 0.50 GPa, which are used in the modals in ANSYS analyzed in Figs. 3 and 4. In these ten zones, *H* is 0.42± 0.04 GPa, 0.47± 0.05 GPa, 0.73± 0.07 GPa, 0.82± 0.10 GPa, 1.75± 0.24 GPa, 0.54± 0.11 GPa, 0.52± 0.06 GPa, 0.79± 0.05 GPa, 0.45± 0.04 GPa and 0.87± 0.19 GPa. *E* and *H* of the veins always have the same trends. Larger values of *E* always appear at the bending location, which is used to unfold the hind wing. *E* in the darker zones of the vein is larger than *E* in the lighter zones. The darker zones of the vein maybe have some special materials to change the nanomechanical properties of the veins for bending.

**Fig. 6.**
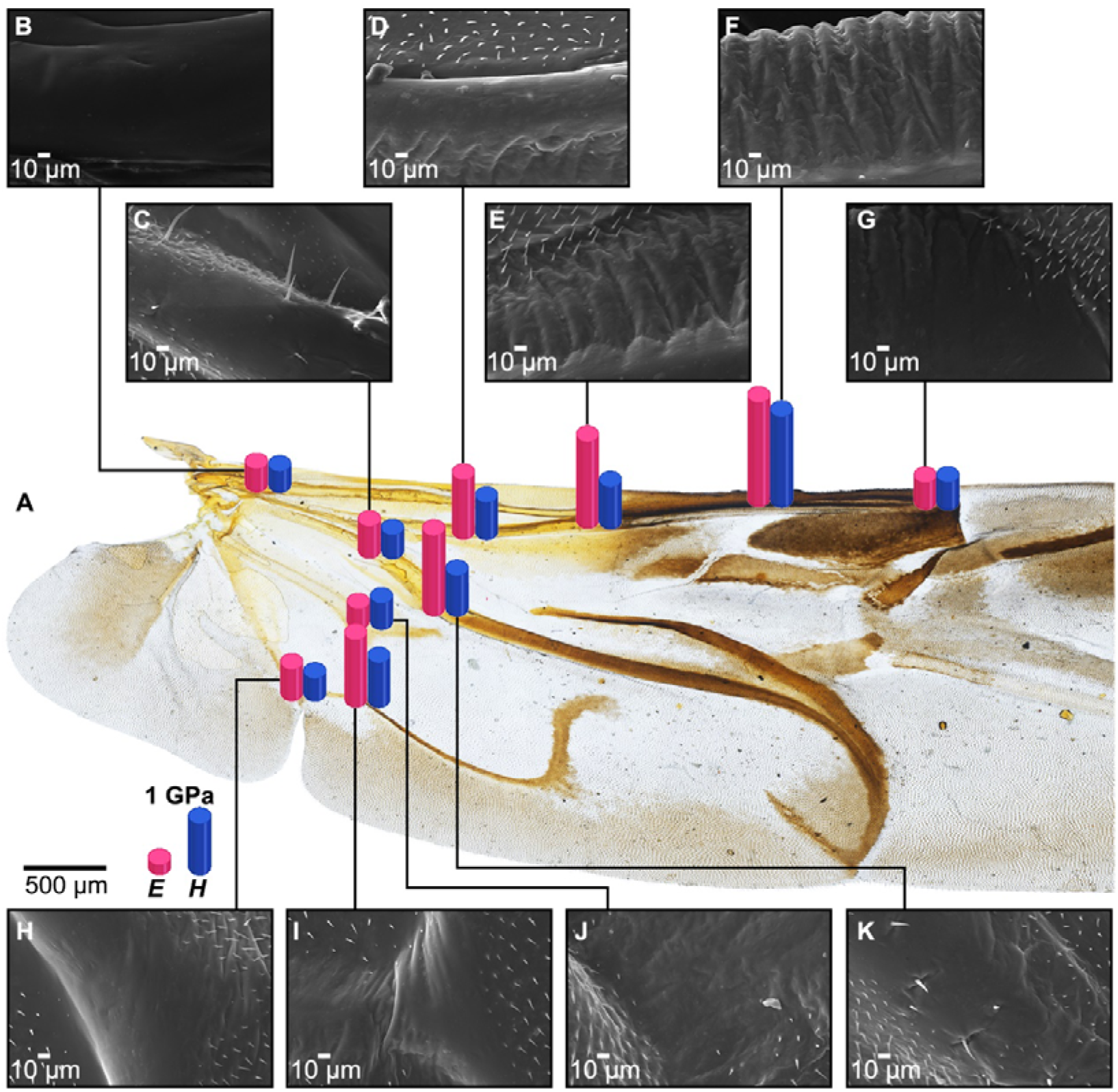
(A) Reduced modulus (*E*) and hardness (*H*) at different positions on the veins, which are investigated by nanoindentation tests. Scale bar: 500 μm and 1 GPa. (B-K) Experiment points in the veins, which are the ten points marked by red hexagons in Fig. 2. Scale bar: 10 μm.

To analyze the nanomechanical properties of the veins of the hind wing using the ANSYS software, different veins with different reduced modulus or the uniform reduced modulus in the two models are considered the same material. The average reduced modulus of the wing membrane is 0.90±0.05 GPa. The average reduced modulus of the veins is 5.17±0.32 GPa. The property settings of the models are based on the nanoindentation measurements.

### 3.4. Mechanical properties of the hind wings

The models of the veins with different reduced modulus are used to compare the displacement of the hind wing with that of the similar models of the veins with the uniform reduced modulus during hovering. As Fig. 7,8 shows, the maximum deformation also appeared at the red regions of the tip of the hind wing, and the minimum deformation is found at the base of the dark blue regions of the hind wing. Furthermore, the base of the wing is constrained so the minimum deformation occurs here. From the base to the apex of the hind wing, the deformation trend of the hind wing is increasing.

**Fig. 7.**
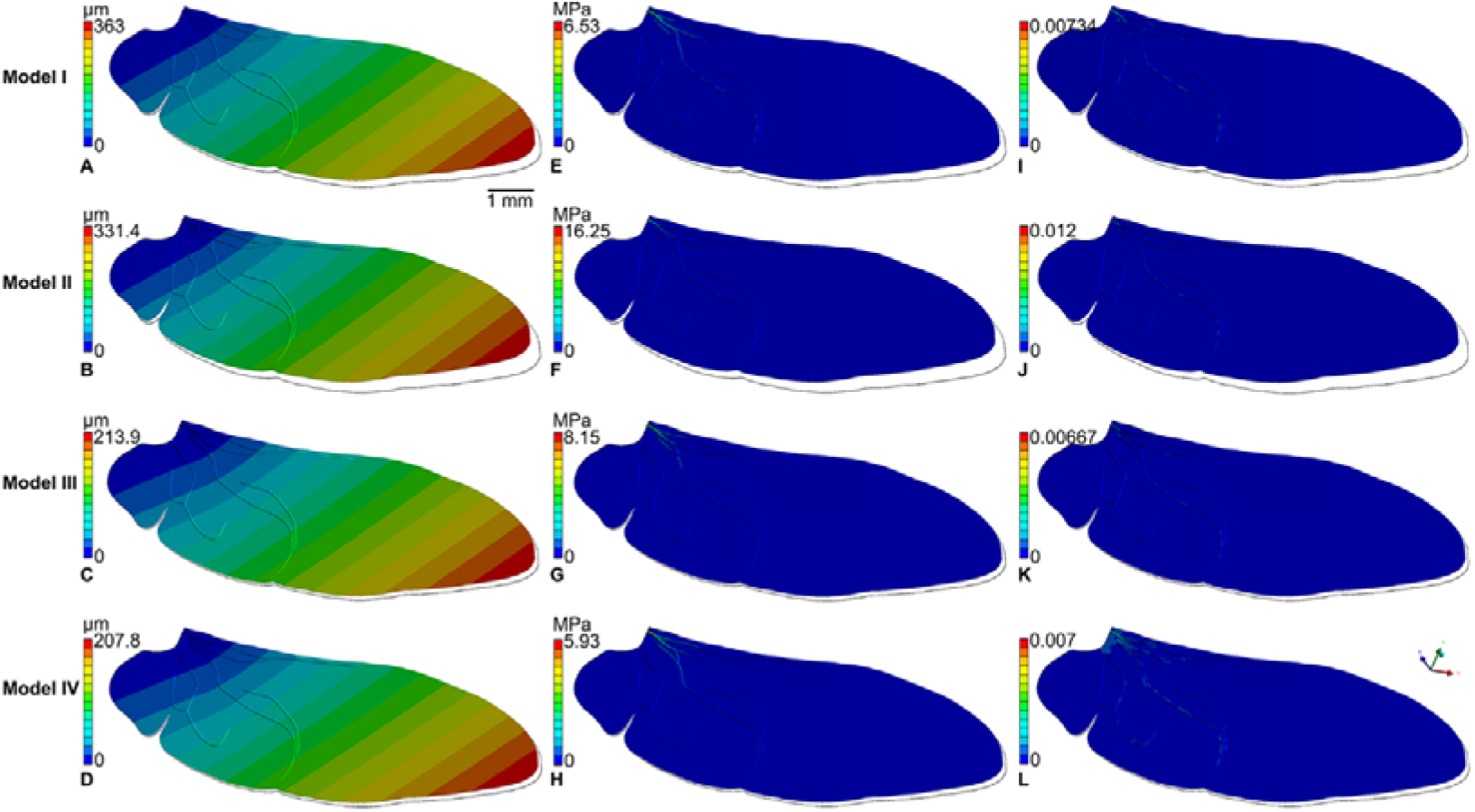
Deformation(A-D), stress(E-H) and elastic strain (I-L) of bending deformation of Model I, Model II, Model III and Model IV.

**Fig. 8.**
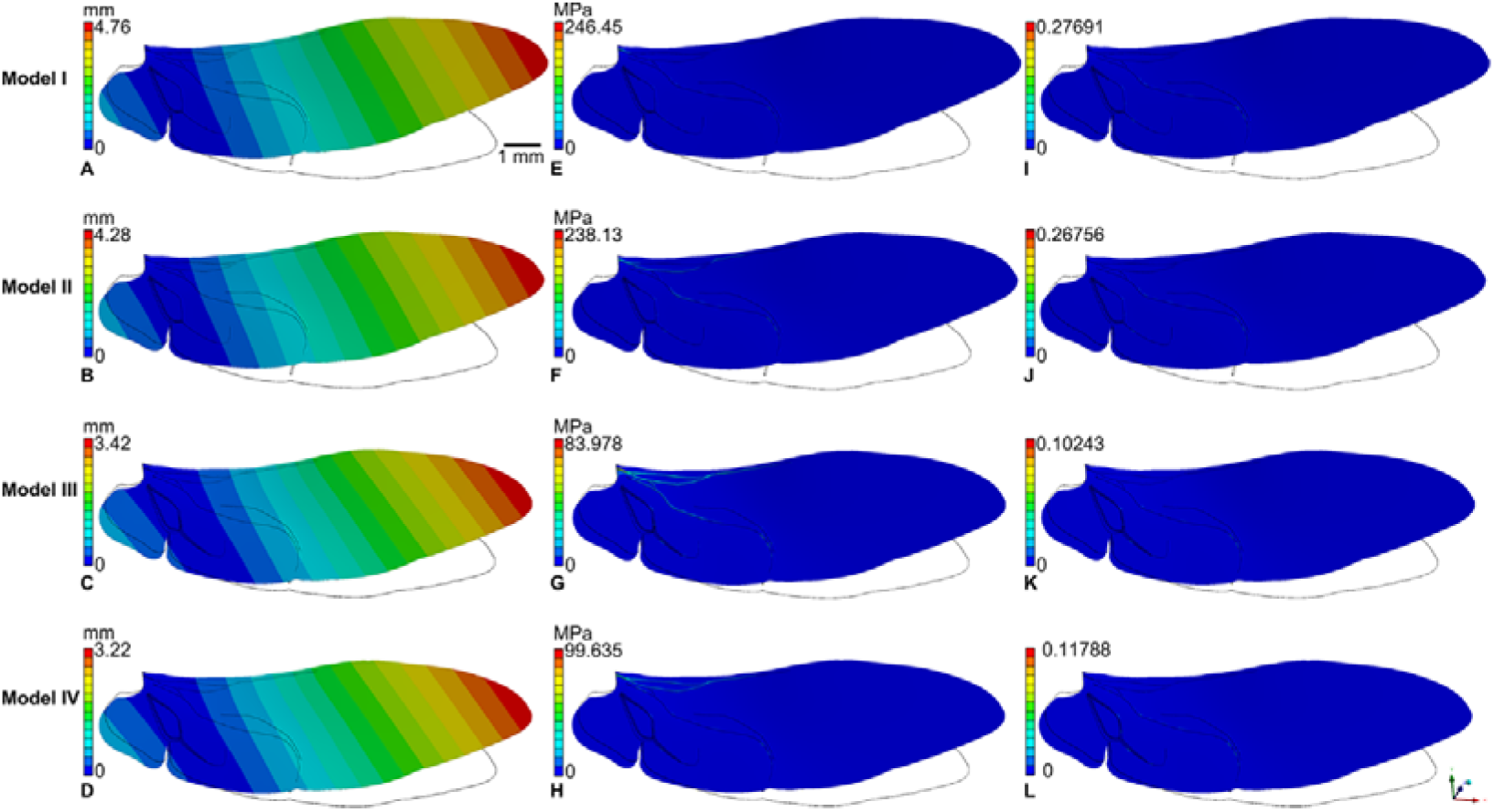
Deformation(A-D), stress(E-H) and elastic strain (I-L) of twisting deformation of Model I, Model II, Model III and Model IV.

Based on the results presented in Fig. 7,8 and results present in Table 1, the hind wing with the uniform reduced-modulus veins undergoes less bending deformation than hind wings with different reduced-modulus veins. However, the bending deformations tendency of the four models are not significantly different. The hind wing models with different and uniform reduced-modulus veins have almost similar twisting deformation tendencies, and the hind wing with different reduced-modulus veins has a larger twisting deformation than those with uniform reduced-modulus veins. The hind wing models with different reduced modulus have a significant effect on the deformation. The bending and twisting deformation of models with wrinkle structure are also larger than the others without wrinkled structure. Thus, the wrinkled structure plays an important role on the flexibility of hind wing as well.

**Table 1.** Maximum numerical analysis results of the bending and twisting deformation of Models I, II, III and IV.

| Model | Bending deformation |  |  | Twisting deformation |  |  |
| --- | --- | --- | --- | --- | --- | --- |
| | Displace /$\mu\text{m}$ | Stress /MPa | Elastic strain | Displace /mm | Stress /MPa | Elastic strain |
| I | 363 | 6.53 | 0.00734 | 4.76 | 246.45 | 0.27691 |
| II | 331.4 | 16.25 | 0.012 | 4.28 | 238.13 | 0.26756 |
| III | 213.9 | 8.15 | 0.00667 | 3.42 | 83.978 | 0.10243 |
| IV | 207.8 | 5.93 | 0.007 | 3.22 | 99.635 | 0.11788 |

Table 1 shows different maximum numerical analysis results for the bending and twisting deformation of Models I, II, III and IV. The four models always have identical trends. Model I has the maximum value, and model IV has the minimum values.

As Fig. 9 shows, the model with real veins with wrinkles has a larger bending deformation than that with veins that are assumed to be smooth circular pipes. The same load of force at the end of the vein is used, 2.3×10^−4^ μN.

**Fig. 9.**
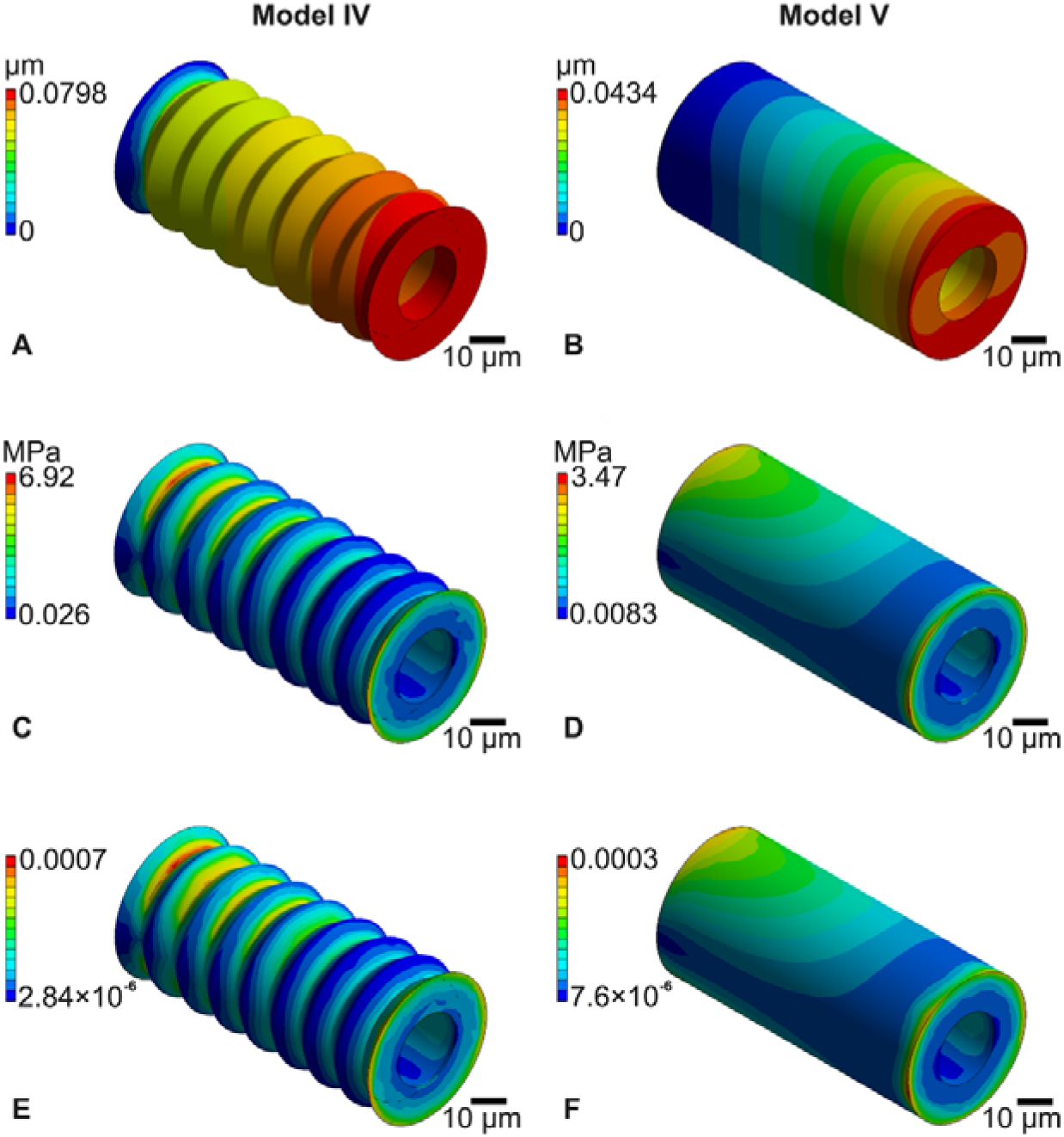
Finite element analysis results of Models IV and V. (A and B) Deformation of the two vein models; (C and D) stress of the two vein models; (E and F) elastic strain of the two vein models.

Based on the results present in Fig. 9, the two vein models also have different bending deformations. We observe that veins with wrinkles have a better deformation tendency and the deformation of the smooth circular pipe is concentrated at the location where force is applied. However, veins with wrinkles have a larger maximum stress than tubular veins. The maximum bending deformations of the vein models with/without wrinkles are 0.08 μm and 0.04 μm, respectively.

## 4. Discussion

In this paper, the hind wing of *H. axyridis* is shown to be a folding/unfolding organ, and bending and twisting deformations, which affect flight performance, result from the nanomechanical properties of different reduced modulus. Importantly, the hind wings can support the body weight and wind resistance during hovering, suggesting the design of the biomimetic wing structure for the deployment of MAVs. This paper investigates the biomechanics of deployable hind wings of the Asian ladybird beetle (*Harmonia axyridis*) and their potential roles in their flight.

### 4.1. Microstructure of the hind wing

The membrane wing thickness, pigmentation and veins form taxon-specific color patterns of the hind wings (Shevtsova et al., 2011). The middle regions of the hind wing with different colors are the most effective regions for the folding/unfolding of the hind wing (Sun et al., 2018b). The cross sections of the hind wing have a corrugated structure, which can change the flight performance (Wang et al., 2017). The main veins can bend, but the false tape spring-like veins can fold and bend and are more prone to twisting when folded or loaded (Betts, 1986b). The tape spring structure allows high-performance bending, folding and twisting and has a smaller mass and general simplicity (Kwok and Pellegrino, 2013). The hind wing is like a tape-spring hinge that can be bent and folded under a load (Mallikarachchi and Pellegrino, 2014). As Fig. 2 shows, the ScP is the windward side during flight and is bent when the wing is folded, so the thicknesses of different positions are different.

Wrinkle veins are also found in the bending zone of beetles (Haas and Beutel, 2001), and wrinkled structures of the veins are always observed on *H. axyridis*. However, the veins in other zones have no wrinkles, and they do not need to bend during the folding progress. The functions of the veins with wrinkles may affect the flight performance and unfolding/folding mechanism of the hind wing.

Different positions of the hind wing have different microtrichia structures. The microtrichia on the hind wing are also used for the interlocking mechanism (Sun et al., 2018a), but the microtrichia on the veins should have another function. The microtrichia may not affect flight performance but may be used for self-cleaning and dust proofing to keep impurities off of the hind wing.

### 4.2. Nanomechanical properties of veins

The properties measured by nanoindentation are informative since previous studies have suggested the importance of passive wing deformation, which is governed by the interactions of the aerodynamics and the structural dynamics of the flapping wings. In our analysis, we considered that the material stiffness of the hind wing (reduced modulus, *E*) is not constant throughout the hind wing in regard to veins to compare the effects of the same *E* value and different *E* values of veins on flight performance. The reduced modulus at the test zone of the ScP increases from the base to the end, but the reduced modulus at the end obviously decreases. The reduced modulus increases along the span of the wing (Herbert et al., 2000b). The reduced modulus is always large at the zone of veins with wrinkles, which is always used for folding the hind wing. The reduced modulus is lowest on smooth veins, and there is little difference from the base to the end of the veins. The reduced modulus and hardness always increase, so the bending zones with wrinkled veins have a high hardness, which may affect the performance of the hind-wing fligh, and can prevent hind wing deformation. The variations around the hind wing of both the reduced modulus and vein structure are of profound interest in the context of the vein’s function.

The reduced modulus varies at different vein zones, and larger modulus are always found in folding zones, which are used to fold the hind wing. The thicknesses of the veins do not affect the numerical reduced modulus of the veins, and the bending function of the veins may be an aspect of this finding. The numerical reduced modulus is linked to the level of bending and vein folding of the hind wings. The veins at the base of the hind wing are tubular, which provides the best resistance to torsion and bending (Betts, 1986b). Because of composites with different reduced-modulus vein structures, the different locations of veins strengthen the bending and wind resistance of the hind wing. The reduced modulus can vary widely within a hind wing (Herbert et al., 2000b), and some proteins, such as resilin, can alter the local properties (Gorb, 1999). The reduced modulus is linked to the level of hind-wing folding, vein bending and the structure of the hind wing. Deformation of the hind wings is limited, so the hind wings are strong, which can help *H. axyridis* in flight. If the veins are sufficiently thick, the wing must use a much larger force to fold the hind wings and *H. axyridis* must consume much more energy, which may be related to the nanomechanical properties.

### 4.3. Mechanical properties of the veins and hind wing

The finite element simulation can replace experimentation to explore the function of veins (Xiang and Du, 2017; Hao and Du, 2018). Alternatively, researchers have developed finite element analysis (FEA)-based models (Chimakurthi et al., 2011) to conveniently analyze the effects of the nanomechanical properties and structure of veins on the deformation of the hind wing. Flexible deformation is involved in the deformation and mechanics of the hindwing. Flexible deformation under different flight modes is a current research hotspot. In each flight mode, the hindwing has different postures, flapping angles, and force conditions.

This paper reports the elastic modulus and hardness of the hind wings of the Asian ladybird beetle (*H. axyridis*) measured by nanoindentation. Computational simulations of the static bending and twisting tests with commercial software (ANSYS) are also presented. Bending and twisting deformations are discussed. As shown in Fig.7, the maximum bending deformation of Model I is nearly 42.75% larger than Model IV. The maximum twisting deformation of Model I is nearly 32.35% larger than Model IV in Fig.8. Model I has the larger twisting and bending deformation, so the flight performance changes during hovering. Model I is compared with the other models with an artificial geometry or modulus. Model III and Model IV are constructed by averaging the modulus of the veins, and we find that this modulus averaging reduced the wing deformation.

The variation ratio of the simulation results based on Model IV which has the lower deformation in Table 2. The hind wing with different reduced modulus a significant change than the one with uniform reduced modulus. And the wrinkled structure plays a role on their deformation as well. But we can find that the effect of reduced modulus is more notable. Thus, the reduced modulus and the wrinkled structure common influence the deformation during flapping flight.

**Table 2:**
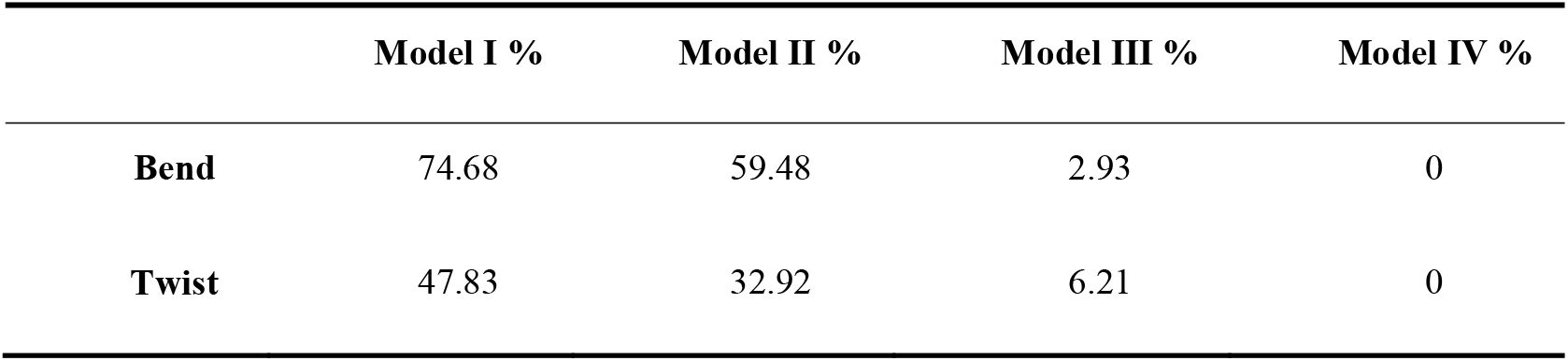
The variation ratio of the simulation results.

Manipulation of the geometry or modulus is an effective method to understand their functions only when the effect of the manipulation on flight performance is appropriately evaluated in detail. The hind wings are always passively deformed to control wing deformations, which can affect the flight performance during flight (Rajabi et al., 2016; Jongerius and Lentink, 2010; Kesel et al., 1998; Mountcastle and Daniel, 2009), so the hind wings has higher flexibility during wind.

Veins with wrinkles are found in the bending zone and allow the hind wing to fold (Haas and Beutel, 2001). As shown in Fig.9, the maximum bending deformation of the vein models with wrinkles is nearly 50% larger than those of the models of veins without wrinkles. The equivalent stress of the vein models with wrinkles is larger than those of the vein models without wrinkles, so veins with wrinkles help to unfold the hind wing. With the same force applied at the end of tube, the two models have different total deformation. The same deformation of the two models illustrates that the value of *F* on the vein with wrinkles is less than that of veins without wrinkles. The same force applied to veins with different structures has different effects, and the maximal deformation of veins with wrinkles is larger than that of veins without wrinkles, so veins with wrinkles also help to fold the hind wing. The deformation of the hind wing is flexible. Macrofurrows must also affect wing stiffness since they should affect the cross-sectional second moment of the area. Veins with wrinkle and various reduced modulus significantly affect the deformation, which changes the flight performance of the hind wing. Flexible deformation of the hind wing has obvious effect on the lifting force. Simple comparisons of the maximum displacement, stress and strains between the realistic and the averaged models under the highly simplified loading conditions are helpful to study the function of veins with wrinkles and to determine the best design for the wing structure of MAVs.

The deformation of wrinkled veins causes the whole vein to bend more except the base zone of the vein, but smooth veins undergo uniform bending in the first half part of the vein, where the force is applied. Therefore, wrinkled veins bend better than smooth veins. This structure is also used for origami designs, which reduces the height and volume of a machine (Morgan et al., 2016). The wrinkles of the ladybird wing-folding mechanism are also further developed into a tubular structure (Gruber, 2008). The wrinkle structure provides more flexibility of passive deformation to control wing deformations. The nanomechanical properties and structure of the veins affect the hind-wing functions of the flight performance and folding mechanism. This wing is a bionic inspiration for the deployable wing structure of MAVs.

To design a wing with a much more efficient flight performance, the veins on the MAV should have different reduced modulus or the uniform reduced modulus. Considering the point of deployable wings of MAVs, the wing structures also use veins with different reduced- modulus to form of an effective structure, so that the deployable wing of MAVs will be much more flexible.

## 5. Conclusion

The wrinkled structures on the veins of the hind wing play an active role in flight and the process of folding/unfolding the hind wing. These structures are favorable for the bending of the hind wings. Special wrinkled structures are found in the bending region of beetle hind wings but not in the wings of most of the other insects that do not fold.

The nanomechanical properties of the veins also affect the flight performance in terms of deformation. A higher hardness mostly corresponds to more deformations, which helps with the wind resistance during beetle flight. However, the structures play more positive roles than the mechanical properties. The wrinkle structures in the bending zone and varying reduced modulus of the veins contribute to the flight performance of bending and twisting deformations of the hind wings.

## Acknowledgements

ZLS and JYS designed the study; ZLS and YWY coordinated the study; ZLS, JT and JYS conducted the research and analyzed the data; ZLS wrote the manuscript; JYS reviewed the manuscript, discussed the results and gave the final approval for publication. Zhiqiang Zhang, ShenYang YuanJie Optics Technology Co.,Ltd, offer technology supporting.

## Data accessibility

This article has no additional data.

## Ethical standard

This work complies with ethical guidelines at Jilin University.

